# Deep mutational scanning of influenza A virus NEP reveals pleiotropic mutations in its N-terminal domain

**DOI:** 10.1101/2024.05.16.594574

**Authors:** Qi Wen Teo, Yiquan Wang, Huibin Lv, Kevin J. Mao, Timothy J.C. Tan, Yang Wei Huan, Joel Rivera-Cardona, Evan K. Shao, Danbi Choi, Zahra Tavakoli Dargani, Christopher B. Brooke, Nicholas C. Wu

**Affiliations:** Department of Biochemistry, University of Illinois at Urbana-Champaign, Urbana, IL 61801, USA; Carl R. Woese Institute for Genomic Biology, University of Illinois at Urbana-Champaign, Urbana, IL 61801, USA; Center for Biophysics and Quantitative Biology, University of Illinois at Urbana-Champaign, Urbana, IL 61801, USA; Department of Microbiology, University of Illinois at Urbana-Champaign, Urbana, IL 61801, USA; Carle Illinois College of Medicine, University of Illinois at Urbana-Champaign, Urbana, IL 61801, USA

**Author notes:** These authors contributed equally to this work (Qi Wen Teo, Yiquan Wang, Huibin Lv). To whom correspondence may be addressed. (N.C.W.).

## Abstract

The influenza A virus nuclear export protein (NEP) is a multifunctional protein that is essential for the viral life cycle and has very high sequence conservation. However, since the open reading frame of NEP largely overlaps with that of another influenza viral protein, non-structural protein 1, it is difficult to infer the functional constraints of NEP based on sequence conservation analysis. Besides, the N-terminal of NEP is structurally disordered, which further complicates the understanding of its function. Here, we systematically measured the replication fitness effects of >1,800 mutations of NEP. Our results show that the N-terminal domain has high mutational tolerance. Additional experiments demonstrate that N-terminal domain mutations pleiotropically affect viral transcription and replication dynamics, host cellular responses, and mammalian adaptation of avian influenza virus. Overall, our study not only advances the functional understanding of NEP, but also provides insights into its evolutionary constraints.

## INTRODUCTION

Influenza A virus (IAV) belongs to the *Orthomyxoviridae* family and is characterized as an enveloped, negative-sense single-stranded, segmented RNA virus. Its genome is comprised of eight viral segments encoding at least 12 proteins^1^. Segment eight, which is the shortest segment, is known as the non-structural (NS) segment. NS segment encodes two proteins via splicing, NS protein 1 (NS1) and nuclear export protein (NEP, also known as NS2)^2^. NS1 is one of the earliest proteins expressed during virus infection and has been well characterized as a potent type I interferon (IFN) antagonist^3^. On the other hand, NEP serves a nuclear export function and is expressed at a later stage during infection as a less abundant spliced product of the NS segment^4^. While IAV with NS1 deletion is replication-competent albeit attenuated^5^, IAV lacking NEP results in a replication-incompetent virus^6^.

NEP has a protease-sensitive N-terminal domain (amino acids 1-53), which harbors two nuclear export signals (NES)^7, 8^, and a protease-resistant C-terminal domain (amino acids 54-121)^9^, which interacts with M1. A major function of NEP is to mediate the nuclear export of influenza viral ribonucleoproteins (vRNPs) for viral packaging^10^. Each influenza vRNP comprises viral RNA (vRNA) binding to three viral polymerase subunits PB2, PB1, and PA, as well as nucleoprotein (NP). NEP orchestrates the nuclear export of vRNPs through its interaction with cellular β-importin protein CRM1^7, 8^ and influenza matrix protein (M1), which in turn binds to the vRNPs^7, 9^. The vRNP complex is responsible for both transcription and replication of the vRNA. Transcription occurs inside the nucleus where the vRNA is transcribed into capped and polyadenylated messenger RNA (mRNA) by a primer-dependent mechanism^11, 12^. For viral genome replication, positive-sense complementary RNA (cRNA) is first produced using vRNA as a template and subsequently serves as a template to produce more vRNA^11, 13^. The synthesis of these three viral RNA species, namely mRNA, cRNA, and vRNA, exhibits distinct dynamics during infection^14, 15^. The timing of these dynamics is important for optimal production of infectious virions^16, 17^. Several studies have suggested that NEP is a polymerase-associated cofactor that acts as a ‘molecular timer’ to regulate the switch from transcription to replication, which is attributed to the gradual accumulation of NEP^17, 18, 19^.

Viral polymerase activity represents a major barrier for avian IAV adaptation to mammalian hosts. Several mammalian adaptive mutations in the polymerase subunits of avian IAV have been identified, as exemplified by E627K in PB2^20^. However, NEP has received increasing attention as a crucial determinant of IAV tropism, facilitating avian IAV to overcome the replication block in mammalian cells^21, 22, 23^. Notably, avian H5N1 polymerase is particularly susceptible to the polymerase-enhancing ability of NEP that harbor mammalian adaptive mutations (e.g., M16I, Y41C, and E75G)^22^. These mammalian adaptive mutations in NEP significantly augment cRNA and vRNA synthesis, as well as mRNA transcription^22^.

While residues 63-116 in the C-terminal domain of NEP form a helical hairpin^9^, the rest of the protein is structurally disordered^9, 24^. This poses a challenge to understand its sequence-function relationship, especially for the N-terminal domain. In this study, the replication fitness effects of ∼2,000 mutations of NEP were measured using deep mutational scanning. Our findings revealed that the N-terminal domain of NEP displays a greater mutational tolerance than its C-terminal domain. Furthermore, we showed that multiple mutations in the N-terminal domain simultaneously affect the transcription-to-replication switch, innate immune response, cellular apoptosis, as well as facilitate mammalian adaptation of avian IAV.

## RESULTS

### Deep mutational scanning of H1N1 A/WSN/1933 NEP

To perform deep mutational scanning of NEP, we utilized the eight-plasmid reverse genetics system of influenza H1N1 A/WSN/1933 (WSN) virus with NS1 and NEP separated by a 2A autoproteolytic cleavage site (**Figure S1A**)^25, 26^. Here, the open reading frames of NS1 and NEP did not overlap, as opposed to NEP being produced as a spliced product of NS1 in the unengineered influenza virus. This virus, herein referred as wild-type NS-split (WT_NS-split_), enabled us to construct a saturation mutagenesis library spanning residues 2-113 of NEP without mutating NS1. Over 98% of the clones in the mutant library contained zero to one amino acid mutation (**Figure S1B**). The virus mutant library was then rescued and passaged once for 24 hours in MDCK-SIAT1 cells. Using next-generation sequencing, the frequencies of individual mutations in the plasmid mutant library and the post-passaged mutant library were measured. Subsequently, the fitness value of each mutation was computed as the normalized log_10_ enrichment ratio such that the fitness value of WT_NS-split_ was 0, whereas beneficial and deleterious mutations would have positive and negative fitness values, respectively. Our deep mutational scanning experiment measured the replication fitness effects of 1,895 (89%) out of 2,128 all possible amino acid mutations across the 112 residues of interest (**Figure 1 and Table S1**). A Pearson correlation of 0.77 was obtained between two biological replicates (**Figure S1C**), demonstrating a reasonable reproducibility of our deep mutational scanning result.

**Figure 1.**
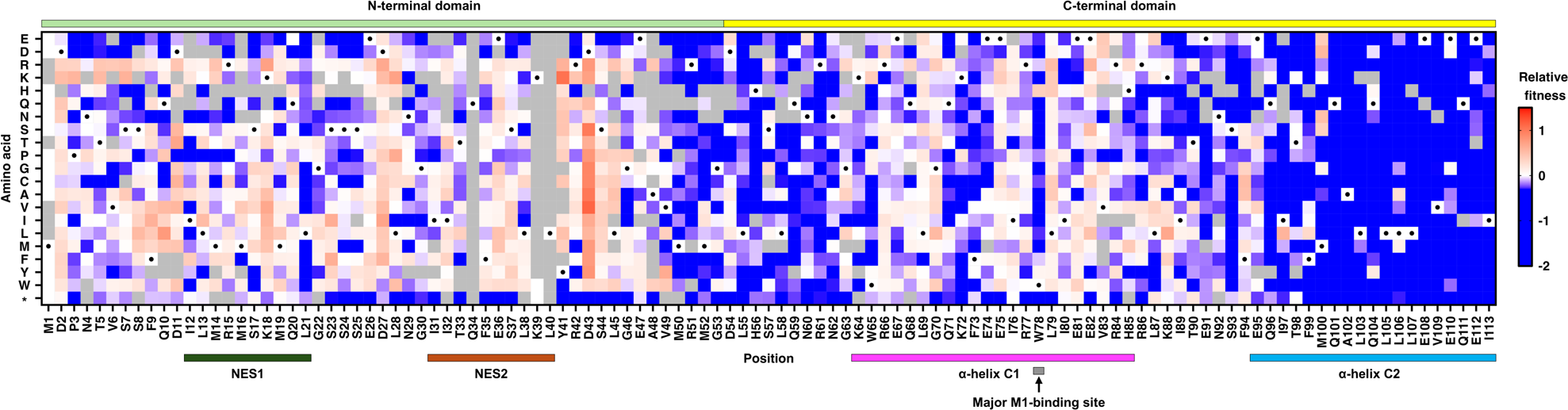
Deep mutational scanning of WSN NEP. The replication fitness effects of individual mutations in the WSN NEP were measured by deep mutational scanning and are shown as a heatmap. Wild-type (WT) amino acids are indicated by a black circle. Mutations in gray were excluded in our data analysis due to low input count. N-terminal domain, C-terminal domain, nuclear export signal (NES), and the M1-binding site are annotated by colored bars. Two α-helices C1 and C2 are annotated by pink and blue bars, respectively. Stop codon is annotated as *.

To further validate our deep mutational scanning result, we constructed seven amino acid mutations individually, including three that had low fitness values (M52F, H56W, and Q59T), and four that had high fitness values (I32T, T33L, S37Y, and D43V). Except residue IIe32, the other residues are evolutionarily highly conserved (**Figure S2**). Subsequently, a virus rescue experiment was performed to assess the replication fitness of these mutations (**Figure 2A**). Consistent with our deep mutational scanning result, mutations I32T, T33L, S37Y, and D43V were rescued to a titer similar to or higher than WT_NS-split_, whereas M52F and Q59T did not produce any detectable infectious particles. Nevertheless, in contrast to the deep mutational scanning result, H56W was rescued to a titer similar to WT_NS-split_. Together, this validation experiment indicates that our deep mutational scanning result contains some measurement errors but is mostly reliable. Of note, as described in a previous study^17^, WT_NS-split_ had a two-log decrease in virus titer compared to the parental WT_non-NS-split_ (**Figure 2A**).

**Figure 2.**
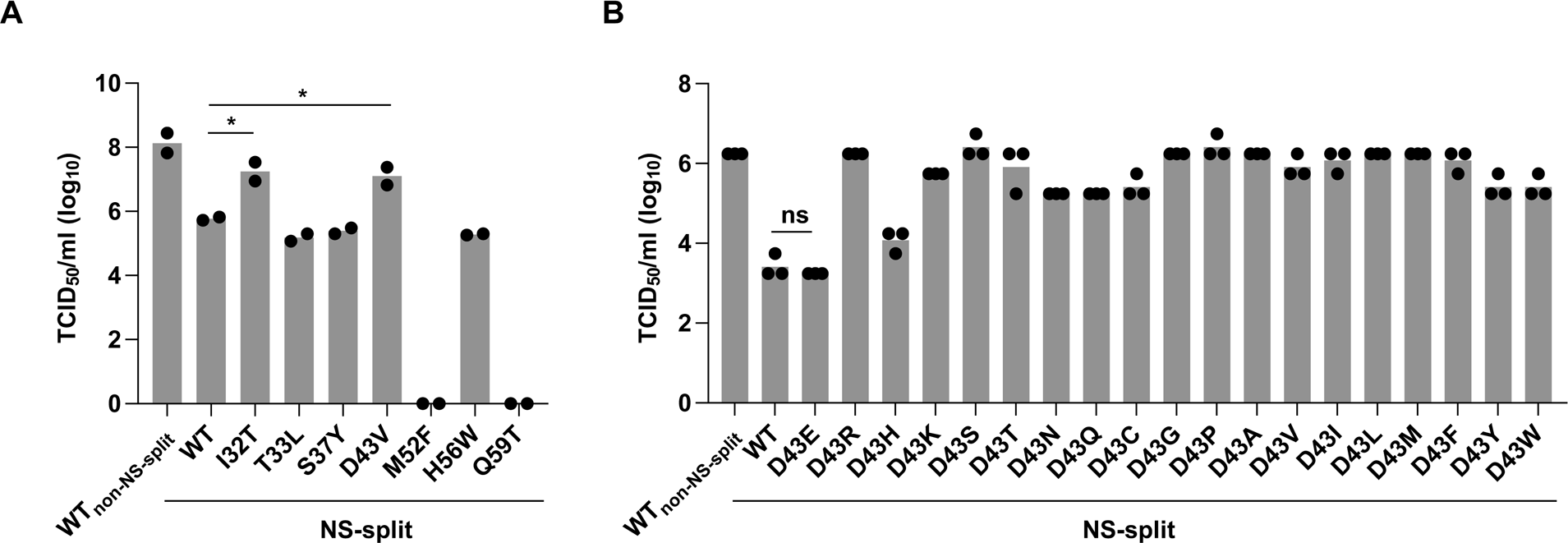
Virus rescue experiment of mutations at residue Asp43. **(A-B)** Mutations were individually introduced into WSN NEP. Their effects on replication fitness were examined by a virus rescue experiment. Virus titer was measured by TCID_50_ assay. **(A)** Replication fitness effects of selected mutations at various residues. Each bar represents the mean of two independent biological replicates. Statistical significance was analyzed by two-tailed Student’s *t*-test: \**P* < 0.05. **(B)** Replication fitness effects of mutations at residue Asp43. Each bar represents the mean of three independent biological replicates. Difference in virus titer between WT_NS-split_ and mutations at residue 43 is statistically significant, except for D43E. Statistical significance was analyzed by two-tailed Student’s *t*-test: ns: not significant. Different batches of cells were used for the virus rescue experiment in (A) and (B).

### Enrichment of mutations with high replication fitness at residue 43

Our deep mutational scanning experiment revealed that the C-terminal domain (average mutational fitness value = -0.36) exhibited lower mutational tolerance than the N-terminal domain (average mutational fitness value = 0, **Figure 1**). This may be partly due to the presence of structural constraints in the C-terminal domain but not the N-terminal domain^9, 24^. The C-terminal domain comprises two α-helices C1 (amino acids 64-85) and C2 (amino acids 94-115), connected by a short interhelical turn (amino acids 86-93)^9^. These helices, largely hydrophobic, form extensive contacts throughout their entire length^9^. The C2 displayed lower mutational tolerance than C1, substantiating previous studies showing that C2 is functionally important in modulating influenza polymerase activity^22, 27^ and regulating the intramolecular interaction with NEP N-terminal domain^23, 28, 29^. Additionally, C2-encoding region is critical for packaging of NS vRNA into the viral particle^30^. Our result also showed that the key M1-binding residue Trp78 in C1 had a low mutational tolerance except for aromatic amino acid mutations (His, Phe, and Tyr), indicating the importance of aromaticity in NEP-M1 interaction.

The N-terminal domain of NEP is featured by two NES (NES1 and NES2)^8, 10^. A previous study has demonstrated that while the NES1 sequence can tolerate some mutations without abolishing its function, certain hydrophobic residues (Met14, Met16, Met19 and Leu21) are crucial^31^. Our data corroborated this finding, revealing that although most mutations within the NES were viable, mutations at hydrophobic residue Leu21 in NES1 were predominantly deleterious (**Figure 1**). In contrast to Leu21, most mutations at residue Asp43, which did not reside in any region with known functions, exhibited high replication fitness in our deep mutational scanning result (**Figure 1**). To experimentally validate this finding, all 19 amino acid mutations were individually introduced into residue 43 of the WT_NS-split_ and analyzed by a virus rescue experiment. Consistent with our deep mutational scanning result, our virus rescue experiment showed that except Glu, which is chemically similar to Asp, all other amino acids mutations increased the titer by 0.5-3 log (**Figure 2B**). In fact, the titer of individual mutants in our virus rescue experiments had a Pearson correlation of 0.78 with the deep mutational scanning result (**Figure S1D**). Collectively, while the N-terminal domain had a high mutational tolerance, certain mutations in the N-terminal domain could dramatically alter virus fitness, highlighting its functional significance.

### N-terminal domain mutations modulate replication dynamics

To delve deeper into the functional relevance of the N-terminal domain in the IAV life cycle, we further characterized those five replication-competent mutations in our first virus rescue experiment (**Figure 2A**). These included I32T, T33L, S37Y, D43V, which resided within the N-terminal domain, and H56W, which was in the structurally disordered region downstream of the N-terminal domain. Specifically, we were interested in understanding how mutations I32T and D43V increased the virus titer (**Figure 2A**). Since NEP has been shown to modulate the production of defective interfering particles (DIPs)^32^, we first investigated the DIP production of different mutants by next-generation sequencing. However, our data indicated that the DIPs could not explain the differential virus production among different mutants (**Figure S3A**). For example, although the titer of I32T_NS-split_ and D43V_NS-split_ was a log higher than WT_NS-split_ in our virus rescue experiment (**Figure 2A**), their DIP production profiles were similar.

A major function of NEP is to regulate switching of viral transcription to replication^16, 17, 18, 33^. Therefore, we hypothesized that the difference in virus replication fitness among these mutants was due to their effects on IAV transcription and replication dynamics. To test our hypothesis, we employed quantitative reverse transcription-PCR (RT-qPCR). Although RT-qPCR is routinely used to probe IAV kinetic profile, it has several drawbacks, including high background and low specificity^34^. Consequently, to complement RT-qPCR, we performed the influenza virus enumerator of RNA transcripts (InVERT) analysis, which is an RNA sequencing (RNA-seq) approach that enables quantification of all three RNA species produced by IAV^14^. Both RT-qPCR and InVERT analyses revealed that the transcription-to-replication switch for WT_NS-split_ occurred earlier than the parental WT_non-NS-split_. Moreover, I32T_NS-split_ and D43V_NS-split_, which had higher replication fitness than WT_NS-split_ (**Figure 2A**), displayed transcription-to-replication switch at a later stage of infection, more closely resembling the parental WT_non-NS-split_ (**Figure 3A and S3B**). Conversely, T33L_NS-split_, which had similar replication fitness as WT_NS-split_ in our rescue experiment (**Figure 2A**), had the switch at an earlier stage of infection (**Figure 3A and S3B**).

**Figure 3.**
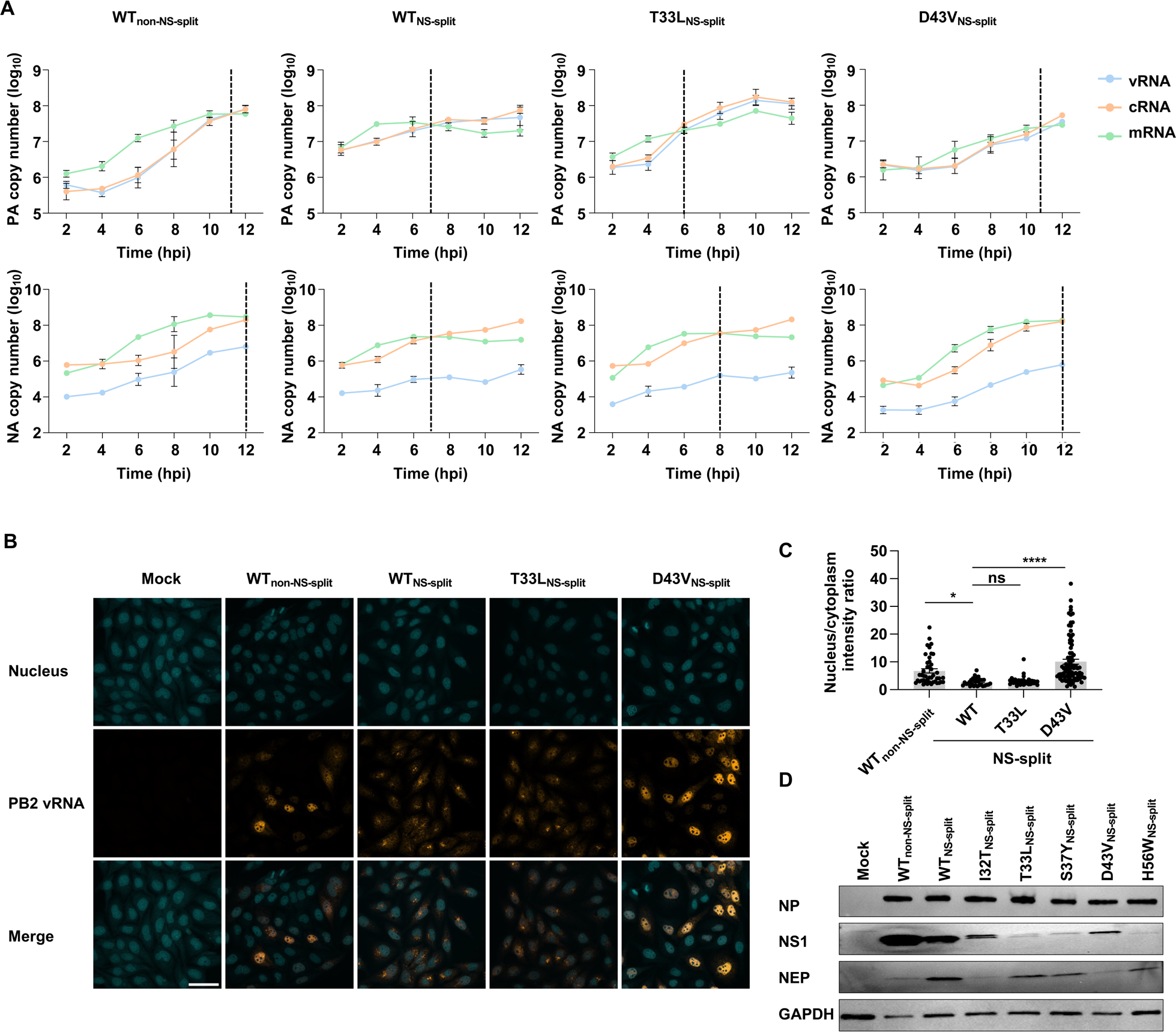
Effects of D43V and T33L mutations on transcription and replication. **(A-D)** MDCK-SIAT1 cells were infected with the indicated mutants at an MOI of 5. **(A)** Kinetics of different IAV RNA species during infection are shown. RNA from infected cells were harvested at the indicated timepoints and analyzed by RT-qPCR. The specificities of the amplified products were confirmed by the melting curve analysis. Data are shown as means of three independent biological replicates. Transcription-to-replication switch was indicated with a vertical dotted line. The switch was defined as the point where mRNA expression intersected with the increasing cRNA expression or reached its plateau level. Blue: vRNA; Orange: cRNA; Green: mRNA. **(B)** Micrographs of FISH staining of the indicated mutants are shown. Infected cells were fixed at 9 hpi, and then stained for PB2 vRNA (orange) and nucleus (cyan). Scale bar: 50 μm. **(C)** Ratio of mean intensity of PB2 vRNA in the nucleus to that of PB2 vRNA in the cytoplasm for micrographs in panel B. Each bar represents the mean ± standard error with individual data points shown (*n* = 46 for WT_non-NS-split_; *n* = 29 for WT_NS-split_; *n* = 39 for T33L_NS-split_; *n* = 91 for D43V_NS-split_). Two outlier data points were removed for cells infected with WT_non-NS-split_, T33L_NS-split_, and D43V_NS-split_, respectively. Statistical significance was analyzed by two-tailed Student’s *t*-test: ns: not significant, \*\**P* < 0.01, and \*\*\*\**P* < 0.0001. **(D)** Western blot analysis of NP, NS1, and NEP protein expression at 16 hpi. GAPDH was used as loading control.

Since NEP facilitates nuclear export of vRNPs^7^, we also performed RNA fluorescence *in situ* hybridization (FISH) to examine if the nuclear export of vRNPs was affected by these mutants. In this experiment, probes that were specific to PB2 vRNA were used. Notably, cells that were infected with WT_NS-split_ and T33L_NS-split_ showed bright perinuclear foci of PB2 vRNA (**Figure 3B and 3C**). This was in stark contrast with the vRNA distribution in cells that were infected with WSN_non-NS-split_ and D43V_NS-split_, where the vRNA was predominantly localized within the nucleus (**Figure 3B and 3C**). These results indicated that the D43V substitution also affected the vRNPs’ nuclear export, despite not being positioned within the NES. Additionally, the timing disparity in the transcription-to-replication switch and the altered vRNPs nuclear export between WT_NS-split_ and D43V_NS-split_ suggested that the increase in replication fitness of D43V may be attributed to altered replication kinetics.

Considering the importance of the NS1 to NEP expression ratio in coordinating the timing of IAV infection^17^, we postulated that the variation of transcription-to-replication switch among mutants correlated with NS expression. Consistently, the parental WT_non-NS-split_ showed a higher NS1 and lower NEP expression compared to WT_NS-split_ (**Figure 3D**). NS1 and NEP expression patterns of mutants with higher replication fitness, namely I32T_NS-split_ and D43V_NS-split_, resembled those of WT_non-NS-split_. By contrast, mutants with lower replication fitness, namely T33L_NS-split_, S37Y_NS-split_, and H56W_NS-split_, showed higher NEP and lower NS1 expressions (**Figure 3D**). These findings suggested that NEP mutations I32T and D43V compensated the fitness defect of WT_NS-split_ by suppressing the NEP expression level to that of WT_non-NS-split_, which may in turn lead to transcription-to-replication switch dynamics akin to that of WT_non-NS-split_.

### Impact of mutations in the N-terminal domain on cellular response regulation

Next, we examined the effects of these mutants on multicycle replication kinetics in both MDCK-SIAT1 and A549 cells. Virus production of I32T_NS-split_ and D43V_NS-split_ was approximately one log higher than WT_NS-split_ at 48 hours post-infection (hpi) in both cell lines (**Figure 4A and S4A**). Notably, although virus production of T33L_NS-split_ was comparable to WT_NS-split_ in MDCK-SIAT1 cell, it was significantly lower than WT_NS-split_ in A549 cells (**Figure 4A and S4A**). The disparity in virus production of T33L_NS-split_ between A549 and MDCK-SIAT1 may be due to differential interferon (IFN) induction in these cell types^35^. These results prompted us to hypothesize that these mutants may affect cellular response during infection, thereby contributing to their different growth kinetics.

**Figure 4.**
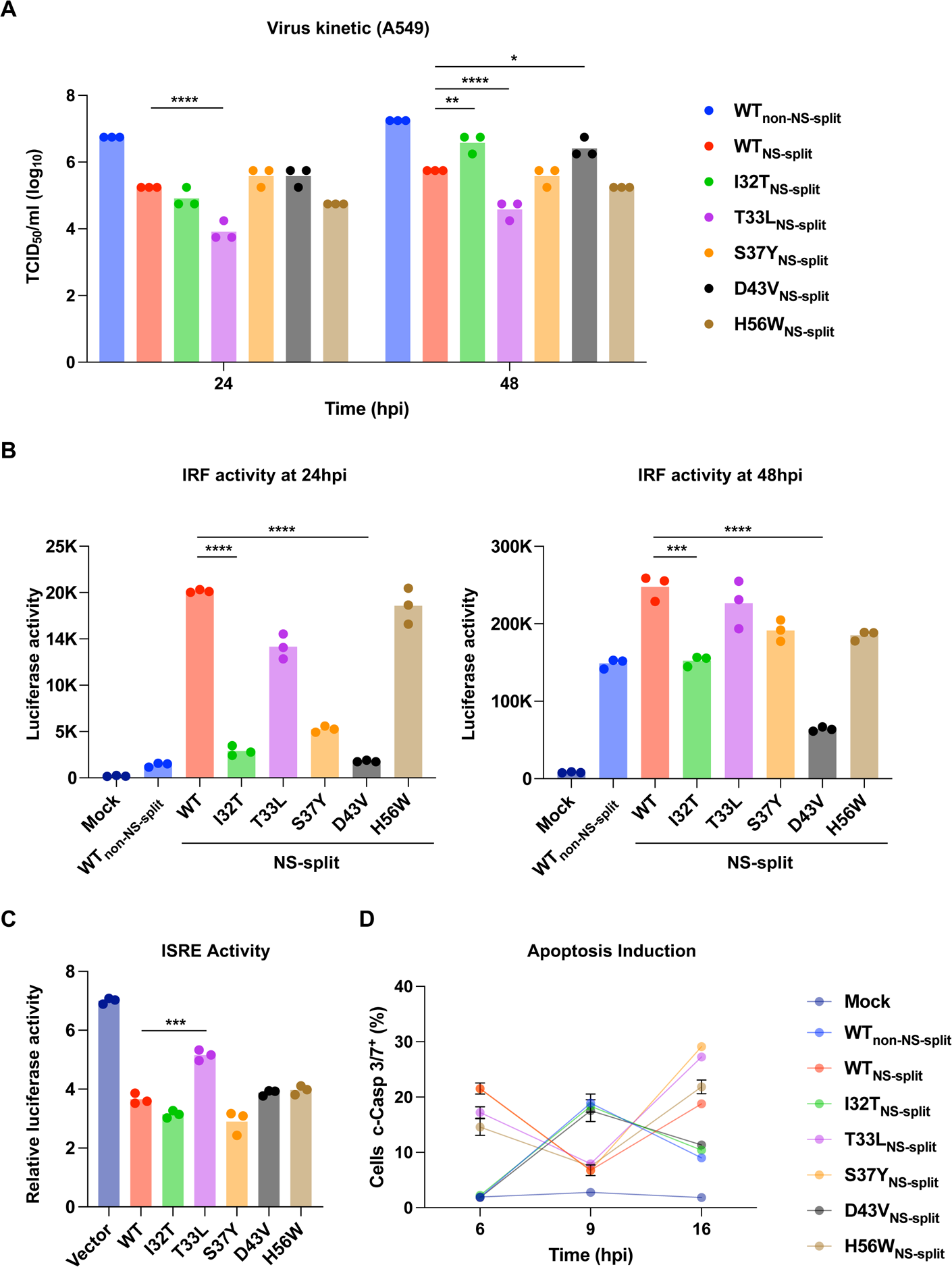
Impacts of NEP mutations on cellular responses. **(A)** A549 cells were infected at an MOI of 0.01 with the indicated mutants. Infectious particles in the supernatant were harvested at the indicated timepoints and quantified via the TCID_50_ assay. Each bar represents the mean of three independent biological replicates. Statistical significance was analyzed by two-way ANOVA: \**P* < 0.05, \*\**P* < 0.01, and \*\*\*\**P* < 0.0001. **(B)** Activation of IRF activity was measured in mock- and IAV-infected A549 cells expressing dual reporters for NF-κB and IRF activities. Each bar represents the mean of three independent biological replicates. Statistical significance was analyzed by Student’s *t*-test: \*\*\**P* < 0.001 and \*\*\*\**P* < 0.0001. **(C)** HEK293T cells were transiently transfected with plasmids expressing the indicated NEP mutants, a plasmid encoding an ISRE-firefly luciferase reporter, and a plasmid expressing Renilla luciferase. At 24 h post-transfection, cells were treated with IFN-I (2000 U/mL) for 16 h prior to measuring luciferase activities. Each bar represents the mean of three independent biological replicates. Statistical significance was analyzed by two-tailed Student’s *t*-test: \*\*\**P* < 0.001. **(D)** A549 cells were infected with the indicated mutants at an MOI of 5. Cells were harvested at the indicated timepoint, and the number of cells undergoing apoptosis were measured using CellEvent Caspase-3/7 Green Flow cytometry kit. Each line represents the mean of three independent biological replicates. Cleaved Caspase-3/7: c-Casp3/7.

A recent study has indicated that NEP promotes innate immunity evasion by interacting with interferon regulatory factor (IRF7) through the N-terminal domain^36^. To ascertain the impact of the NEP mutants on innate immunity, we infected A549^Dual^ cells which allow simultaneously monitoring of IRF activity and NF-κB induction^37^. Our data demonstrated that I32T_NS-split_ and D43V_NS-split_ had significantly lower IRF and NF-κB induction compared to the WT_NS-split_ at 24 and 48 hpi (**Figure 4B and S4B**). This observation helps explain the higher virus production in I32T_NS-split_ and D43V_NS-split_ compared to the WT_NS-split_ at 48 hpi (**Figure 4A and S4A**). NEP has also been shown to suppress ISRE activation^36^. As a result, we further tested if these NEP mutants affect ISRE activity by transiently transfecting HEK293T cells with NEP expression constructs, an ISRE-firefly luciferase reporter plasmid, and a constitutively expressed Renilla luciferase plasmid for normalization. Consistently, we showed that NEP suppressed ISRE activity in response to recombinant IFN-α (**Figure 4C**). This inhibition can be observed across all NEP mutants with varying protein expression levels (**Figure 4C and S4C**). Our result also indicated that T33L mutation weakened the ability of NEP to antagonize IFN response despite comparable, if not higher, protein expression level with other mutants (**Figure 4C**). This helps explain the lower virus production observed in T33L_NS-split_ compared to that of WT_NS-split_ in A549 cells (**Figure 4A**).

When the innate immune responses fail to effectively control IAV replication, cells may activate a secondary antiviral response via programmed death, known as apoptosis^38^. However, IAV strategically inhibits early induction of apoptosis to prolong replication and induces apoptosis at later stages of infection to enhance viral spread^39, 40, 41, 42^. Given the difference in innate immune response among mutants and unexpected extensive cell death observed in some mutants (WT_NS-split_, T33L_NS-split_, S37Y_NS-split_, and H56W_NS-split_) (**Figure S4D**), we next examined the apoptosis pathway in infected A549 cells. Our results revealed that while the parental WT_non-NS-split_, I32T_NS-split_ and D43V_NS-split_ induced apoptosis at 9 hpi, WT_NS-split_, T33L_NS-split_, S37Y_NS-split_, and H56W_NS-split_ infection resulted in biphasic induction of apoptosis, occurring as early as 6 hpi and again at 16 hpi **(Figure 4D**). Therefore, the differential viral RNA transcription-to-replication switch among these mutants may not solely account for their difference in replication fitness due to the intricate interplay between virus and cellular responses. Taken together, our findings suggest that the N-terminal domain of NEP modulates virus production through regulating multiple cellular responses.

### NEP mutations increase the polymerase activity of avian IAV in human cells

Previous studies have shown that mutations in the C-terminal domain of NEP can modulate the polymerase activity^23, 43^. In this study, we sought to explore whether the N-terminal domain also influenced polymerase activity. We first performed an experiment by testing the effects of our NEP mutants on the WSN polymerase, using a polymerase reconstitution assay in HEK293T cells. Nevertheless, no discernible differences in WSN polymerase activity were observed across all tested NEP mutants, irrespective of the amount of NEP present (**Figure 5A**).

**Figure 5.**
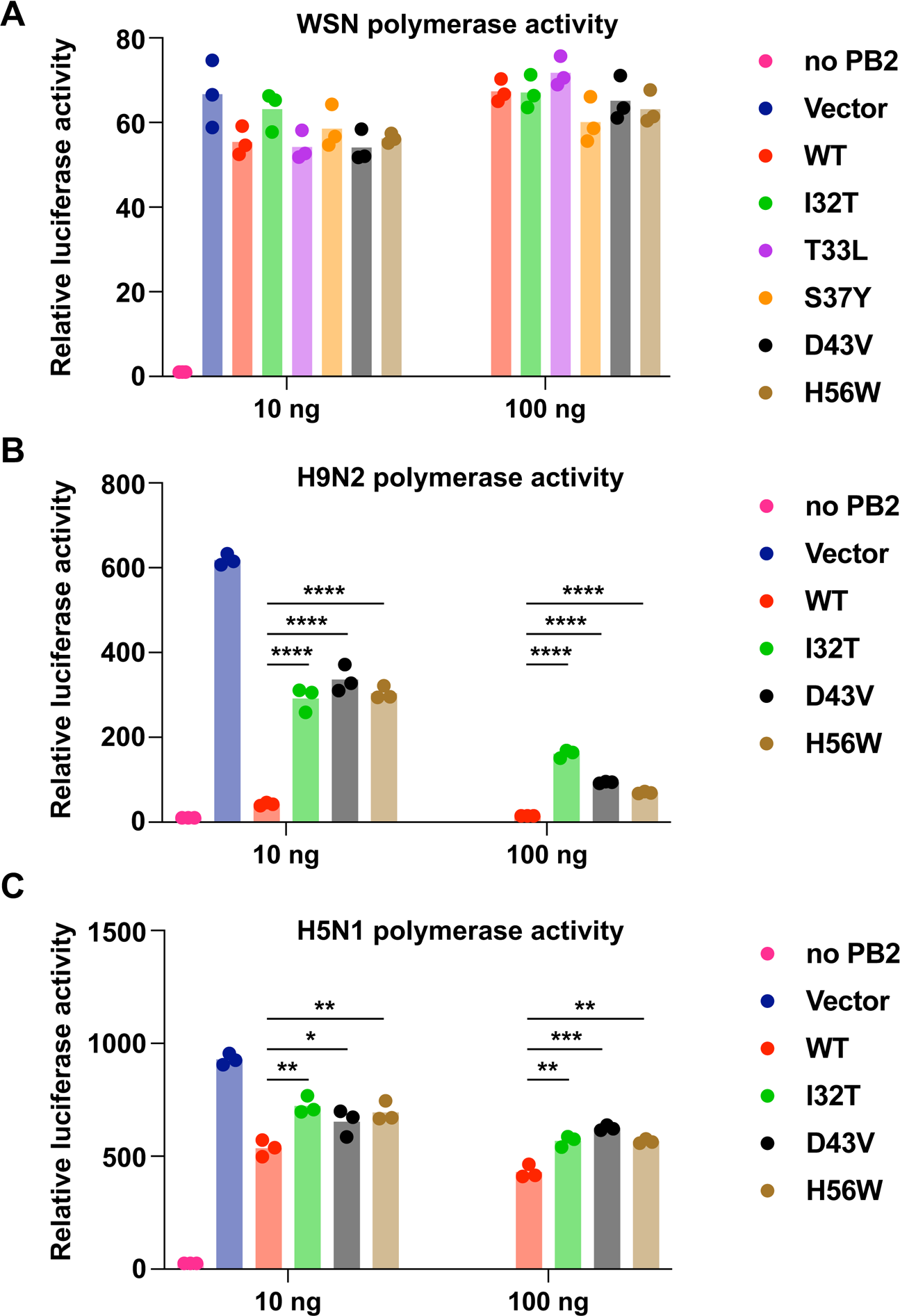
Polymerase activity of avian influenza in cultured human cells. Influenza polymerase reconstitution assay in HEK293T cells for **(A)** WSN, **(B)** A/Quail/Hong Kong/G1/97 (H9N2) and **(C)** A/Vietnam/1203/2004 (H5N1). HEK293T cells were transiently transfected with plasmids encoding the RNP complexes (PB1, PB2, PA, and NP) and vector or indicated amount of plasmids encoding NEP mutants, together with human polymerase I-driven vRNA and Renilla-expressing luciferase reporters. Renilla activity was used to normalize variation in transfection efficiency. Measurements were taken at 16 h post-transfection. Each bar represents the mean of three independent biological replicates. Statistical significance was analyzed by Student’s *t*-test: \**P* < 0.05, \*\**P* < 0.01, \*\*\**P* < 0.001 and \*\*\*\**P* < 0.0001.

However, given that there is growing evidence indicating that mammalian-adaptative mutations in NEP enhance avian IAV replication in cultured human cells^21, 22, 23, 44, 45^, we aimed to further investigate the effect of these NEP mutants on the avian IAV polymerase activity in human cells. We individually introduced each of the mutations I32T, D43V, and H56W into NEP from a low pathogenic avian influenza (LPAI) strain, A/Quail/Hong Kong/G1/97 (H9N2), and a highly pathogenic avian influenza (HPAI) strain, A/Vietnam/1203/2004 (H5N1). Subsequently, we assessed their polymerase activity in HEK293T cells. Our data revealed that all three NEP mutants partially increased the polymerase activity of the H9N2 and H5N1 strains (**Figure 5B and 5C**). Surprisingly, H56W mutant also enhanced the avian polymerase activity (**Figure 5B and 5C**), despite having very different phenotypes compared to I32T and D43V mutants in terms of protein expression level (**Figure 3D and S3C**), IRF induction (**Figure 4B**), and apoptosis induction (**Figure 4D**). In sum, these findings suggest that I32T, D43V and H56W mutations in the NEP of H9N2 and H5N1 can potentially facilitate mammalian adaptation of avian IAV.

## DISCUSSION

For years, NEP has been perceived as a non-structural protein when purified from virions^46, 47^. It was not until 1988 that the nuclear export function of NEP was unveiled^7^, which represented a pivotal discovery in influenza virus biology. As our understanding of NEP has evolved, its multifaceted roles have emerged as focal points of investigation. However, more than 25 years since the discovery of its nuclear export function, the sequence-function relationships of NEP remain largely obscure. By systematically defining the mutational fitness landscape of influenza NEP, this study advances our understanding of its functional constraints.

A highlight of the study is that mutations at N-terminal domain of NEP display pleiotropic effects including viral RNA synthesis dynamics, innate immune response modulation, apoptosis induction, and mammalian-adaptation of avian IAV. We speculate that the pleiotropic phenotypes of the N-terminal domain mutations stem from alterations in the NEP:NS1 expression ratio, which subsequently perturbs viral RNA synthesis dynamics and cellular responses. Nonetheless, the modulation of apoptosis by NEP mutations was not expected. While NS1 is known to be an important regulator of host apoptotic cell death^39, 41^, the role of NEP in apoptosis is less clear. Although our study shows that the NEP mutant viruses induce monophasic or biphasic apoptosis, it is possible that the NS1 expression level is the primary leading cause of this phenotype. Future investigation is required to dissect the underlying mechanisms.

One interesting aspect of overlapping open reading frames is how nucleotide usage is balanced to according to the functions of each open reading frame^48^. A previous study has found that the evolution of the overlapping regions between Tat and Rev of HIV-1 is constrained by the function of either but not both of the proteins^48^. In the case of NS protein, majority of the N-terminal domain of NEP overlaps with the C-terminal effector domain of NS1, which predominantly suppresses the host response and is associated with the pathogenicity of the virus^49, 50^. Given the high mutational tolerance in the disordered N-terminal of NEP and the role of NS1 as a key immunomodulatory factor, it is likely that NEP accommodates the nucleotide preferences of NS1 due to functional constrains imposed by NS1. In other words, the sequence conservation of the N-terminal domain is largely due to the overlapping open reading frames with NS1, rather than its own functional constraints. Of note, overlapping open reading frames of NS proteins necessitates the use of the ‘NS-split’ system in the present study. However, this hampers our understanding of how mutations within this domain would impact virus replication kinetics as well as cellular responses in natural infection.

Most reported mammalian-adaptive mutations in avian IAV NEP occur in the N-terminal domain^22, 23^, although previous studies have highlighted the importance of the three amino acid at the C-terminal end of NEP for its polymerase-enhancing function^23, 43^. Additionally, mutation T48N in the N-terminal domain of NEP from pandemic H1N1/2009-like recombinant virus (rH1N1pdm) has been found to enhance viral replication in guinea pigs^51^, further consolidating our hypothesis that N-terminal domain of NEP is an important tropism determinant. One notable observation in this study is that all tested mutations in the N-terminal domain of NEP were found to enhance the polymerase activity of avian IAV but not WSN in mammalian cells. A previous study has suggested that NEP has two distinct conformations, “open” and “closed”, and that mammalian adaptive mutations in NEP shift the equilibrium towards the “open” conformation^23^. In our study, we used the lab-adapted A/WSN/33 strain, which is known for robust replicates in mammalian cell culture. Therefore, WSN NEP may already favor the “open” conformation without additional mammalian adaptive mutations, although this speculation requires detailed investigations in the future. Additionally, whether mammalian-adaptive mutations in NEP influence its interaction with the vRNP, thus leading to enhanced polymerase activity, remains an unanswered question.

## ACKNOWLEDGEMENTS

We thank the Roy J. Carver Biotechnology Center at the University of Illinois at Urbana-Champaign for assistance with next-generation sequencing and RNA sequencing. We are grateful to the Institute of Genomic Biology Core Facility at the University of Illinois at Urbana-Champaign for providing access to the LSM880 microscope system. This work was supported by the Searle Scholars Program (N.C.W.) and National Institutes of Health (NIH) R01 AI165475 (N.C.W.) and R01 AI139246 (C.B.B.).

## AUTHOR CONTRIBUTIONS

Q.W.T., Y.W., H.L., and N.C.W. conceived and designed the study. Y.W. constructed the NEP deep mutational scanning library and prepared the sequencing libraries. Q.W.T. prepared the samples for RNA sequencing. Q.W.T., Y.W., and N.C.W. performed data analysis. Q.W.T., H.L., K.J.M., Y.W.H., T.J.C.T., J.R.-C., E.K.S., D.C., Z.T.D., and C.B.B. assisted with experiments. Q.W.T., Y.W., H.L., and N.C.W. wrote the paper and all authors reviewed and/or edited the paper.

## DECLARATION OF INTERESTS

N.C.W. consults for HeliXon. The authors declare no other competing interests.

## METHODS

### Cells and viruses

Madin-Darby canine kidney cells with stable expression of human α-2,6-sialyltransferase (MDCK-SIAT1) (Sigma, catalog #: 05071502-1VL), human lung adenocarcinoma epithelial A549-Dual cells (InvivoGen, catalog #: a549-nfis) and human embryonic kidney HEK293T (ATCC, catalog #: CRL-3216) cells were maintained in Dulbecco’s modified Eagle’s medium (DMEM; Gibco) supplemented with 10% fetal bovine serum (FBS; Gibco), 100 U/mL Penicillin-Streptomycin. Cells were cultured in humidified incubators at 37 °C with 5% CO_2_. Recombinant influenza A/WSN/1933 (H1N1/WSN) virus was generated using the pHW2000 eight-plasmid reverse genetics system^26^. Viral genotypes were confirmed by sequencing vial genomes.

### Mutant library construction

The mutant library was constructed based on the pHW2000 eight-plasmid reverse genetics system for influenza virus. The linearized vector and a library of mutant WSN NEP inserts were generated through PCR. The linearized vector was generated using 5’-CGA CCG TCT CTG GGG CCG GGA GGT CGC GTC ACC GAC-3’ and 5’-CGA CCG TCT CTT CTC CTC GAC GTC CCC GGC TTG CTT-3’ as primers. For insert generation, two batches of PCRs were performed, followed by overlapping PCRs. The first batch of PCRs consisted of 15 reactions, each with an equal molar mix of eight forward primers, as the forward primer and a universal reverse primer 5’-CGA CCG TCT CTC CCC GGG GGA GGT ATA TCT TT-3’. The forward primers for the first batch of PCRs are listed in Table S2. These forward primers were named as cassetteX_N, in which X represents the cassette ID and N represents the primer number. Forward primers with the same cassette ID were mixed at equal molar ratios and used in the same single PCR. The second batch of PCRs consisted of another 15 reactions, each with a universal forward primer 5’-CGT TAC CCG GCC AAT GCA CGT TTA CGC CAC AAA TTT CTC TCT CCT CAA GCA AGC CGG GGA CGT CTC GGA GAA TCC CGG GCC C-3’. and unique reverse primers as listed in Table S2. Subsequently, 15 overlapping PCRs were performed using the universal forward primer and reverse primer. For each overlapping PCR, the template was a mixture of 10 ng each of the corresponding products from the first and second batches of PCRs. The complete insert was an equal molar mix of the products of these 15 overlapping PCRs. All PCRs were performed using PrimeSTAR Max polymerase (Takara Bio) according to the manufacturer’s instructions. PCR products were purified using Monarch DNA Gel Extraction Kit (New England Biolabs). Both the vector and the complete insert were digested by BsmBI (New England Biolabs) and ligated using T4 DNA ligase (New England Biolabs). The ligated product was transformed into MegaX DH10B T1R cells (Thermo Fisher Scientific). At least one million colonies were collected. Plasmid mutant libraries were purified from the bacteria colonies using PureLink HiPure Expi Plasmid Purification Kit (Thermo Fisher Scientific).

### Rescuing, passaging and sequencing the mutant library

The virus mutant library was rescued by transfecting a co-culture of HEK293T and MDCK-SIAT1 cells (ratio of 6:1) at 60% confluence in a T75 flask. Transfection was performed using Lipofectamine 2000 (Thermo Fisher Scientific) according to the manufacturer’s instructions. At 24 h post-transfection, cells were washed twice with PBS and cell culture medium was replaced with OPTI-MEM medium supplemented with 0.8 μg/ml TPCK-trypsin. The virus mutant library was harvested at 72 h post-transfection, titered by TCID_50_ assay using MDCK-SIAT1 cells then stored at –80°C until use. To passage the virus mutant libraries, MDCK-SIAT1 cells in a T75 flask were washed twice with PBS and then infected with a multiplicity of infection (MOI) of 0.02 in OPTI-MEM medium supplemented with 0.8 μg/ml TPCK-trypsin. At 2 h post-infection, infected cells were washed twice with PBS and fresh OPTI-MEM medium supplemented with 0.8 μg/ml TPCK-trypsin was added to the cells. At 24 h post-infection, supernatant was harvested. Each replicate was transfected and passaged independently.

Viral RNA of the post-passaged mutant library was extracted using QIAamp Viral RNA Mini Kit (QIAGEN). The extracted RNA was then reverse transcribed to cDNA using Superscript III reverse transcriptase (Thermo Fisher Scientific). To add part of the adapter sequence required for Illumina sequencing, the plasmid mutant library and the cDNA from the post-infection viral mutant library were amplified by PCR using primers: 5’-ACT CTT TCC CTA CAC GAC GCT CTT CCG ATC TNN NNN NNN GAC GTC GAG GAG AAT CCC GGG -3’ and 5’-ACT GGA GTT CAG ACG TGT GCT CTT CCG ATC TNN NNG TAG AAA CAA GGG TGT TTT TTA TTA-3’. A total of 12 N were included in each amplicon product as unique molecular identifiers (UMIs) to distinguish true mutations from sequencing errors^52, 53, 54^. For each sample, the six amplicon PCR products were mixed at equal molar ratio. Subsequently, 1 million copies of mixed amplicon PCR products were used as template for a second PCR to add the rest of the adapter sequence and index to the amplicon using primers: 5’-AAT GAT ACG GCG ACC ACC GAG ATC TAC ACX XXX XXX XAC ACT CTT TCC CTA CAC GAC GCT-3’, and 5’-CAA GCA GAA GAC GGC ATA CGA GAT XXX XXX XXG TGA CTG GAG TTC AGA CGT GTG CT-3’. Positions annotated by an “X” represented the nucleotides for the index sequence. The final PCR products were purified by PureLink PCR purification kit (Thermo Fisher Scientific) and submitted for next-generation sequencing using Illumina MiSeq PE250.

### Analysis of sequencing data for deep mutational scanning

Sequencing data was obtained in FASTQ format and analyzed using a custom snakemake pipeline^55^. Firstly, paired-end reads with the same Unique Molecular Identifiers (UMIs) were merged using a Python script. A consensus sequence was created for UMIs with a minimum of two identical sequences. Groups were retained if at least 70% of the sequences agreed on the consensus; otherwise, the group was discarded. Next, primer sequences were removed using cutadapt^56^, followed by the merging of forward and reverse consensus reads using FLASH^57^. The resulting merged consensus reads were processed using the SeqIO module in BioPython^58^, translated into amino acid sequences, and filtered based on their sequence length matching the reference amplicon. Subsequently, mutations were identified by comparing the amino acid sequences to the reference, and merged consensus reads with more than one amino acid mutation were excluded from downstream analysis. The frequency of mutant i in individual samples was normalized as follows:

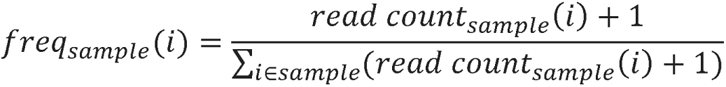

For each mutant *i*, the replication fitness was calculated as follows:

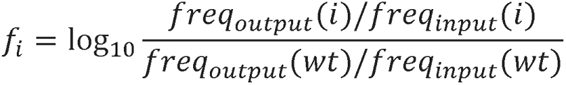

where the *freq_output_* (*i*) represents the normalized frequency corresponding to variant i in the post-passaging virus mutant library, and the *freq_input_* (*i*) represents the normalized enrichment corresponding to variant i in the plasmid mutant library.

### RNA library preparation and sequencing

MDCK-SIAT1 cells were infected with viruses at an MOI of 0.5 and incubated at 4 °C for 30 min to allow the virus to adsorb to the cell surface but not to be endocytosed. Synchronized IAV infection was initiated by shifting the temperature to 37 °C. Secondary infection was blocked using 20 mM NH_4_CI. Cells were harvested at 0, 2, 4, 6, 8, and 12 h post-temperature shift for RNA sequencing to quantify the cellular RNA and viral RNA levels. Uninfected MDCK-SIAT1 cells were also collected for a negative control. RNA from the cells were extracted using RNeasy Plus Micro Kit (Qiagen). To assay both positive- and negative-sense RNAs, total RNA was prepped using the Clontech Pico SMARTer stranded total RNA-Seq kit v2, which preserved strand orientation of the RNA sequences by the template-switching reactions. The cDNA library was then subjected to next-generation sequencing using two lanes of Illumina NovaSeq X 10B PE150. At least 20 million paired-end reads were obtained per sample.

### Quantification of different influenza RNA species

Raw reads underwent adapter trimming using cutadapt^56^. Trimmed reads were then aligned to the reference genome using STAR^59^. The expression levels of all viral genes were initially quantified using htseq-count^60^ to count reads that were assigned to different features in the annotation file and calculate fragments per kilobase of transcript per million mapped reads (FPKM) for each feature. Distinction between viral negative-sense RNA (vRNA) and positive-sense RNA (cRNA and mRNA) was achieved by providing an annotation file specifying RNA strandedness during quantification. cRNA and mRNA were differentiated based on the sequence following five adenosines (5A) at the 3’ end of the gene. During mRNA transcription, the 5A sequence extends to form the poly(A) tail. In contrast, all cRNAs contain a 16-nt sequence at the 3’ end, 13 of which are conserved across segments following the 5A. For each segment in every sample, the pipeline counted sequencing reads that contained the 5A and at least two subsequent nucleotides with complete alignment to the 16-nt consensus sequence as cRNA reads. Reads that contained more than two additional adenosines after the 5A are identified as mRNA reads. The expression values of cRNA and mRNA were determined by multiplying the total read count of positive-sense RNAs.

### Sequencing analysis of defective interfering particles (DIPs)

Raw sequencing reads were fed into our RNA species pipeline for viral negative-sense RNA (vRNA) quantification. Following alignment to the viral genome, identification and quantification of vRNA containing deletion junctions were performed using a custom Python script. To account for the variation of sequencing read coverage between samples, DIPs percentage was derived by normalizing the deletion junction count to the coverage depth of the corresponding vRNA segments.

### Reverse transcription and quantitative PCR

MDCK-SIAT1 cells were infected with viruses at an MOI of 0.5 and incubated at 4 °C for 30 min to allow the virus to adsorb to the cell surface but not to be endocytosed. Synchronized IAV infection was initiated by shifting the temperature to 37 °C. Cells were harvested at 0, 2, 4, 6, 8, and 12 h post-temperature shift. Uninfected MDCK-SIAT1 cells were also collected as a negative control. Total cellular RNA was isolated using RNeasy Mini Kit (Qiagen). Reverse transcription (RT) of RNA was performed using ProtoScript II First Strand cDNA Synthesis Kit (New England Biolabs), in accordance with the manufacturer’s manual. Oligo-dT primer was used in RT reaction for detection of mRNAs, whereas uni-12- (5’-AGC AAA AGC AGG-3’) and uni-13- (5’-AGT AGA AAC AAG G-3’) specific primers were used for vRNAs and cRNA, respectively^61, 62^. Quantification PCR (qPCR) mixtures were prepared according to the user manual of iTaq Universal SYBR Green Supermix (Bio-Rad) and reactions were run in a CFX Opus 96 Real-Time PCR Instrument (Bio-Rad). Segment-specific primers were used for the qPCR analysis. WSN PA forward primer: 5’-CTG ACC CAA GAC TTG AAC CAC-3’; WSN PA reverse primer: 5’-AGC ATA TCT CCT ATC TCA AGA ACA-3’; WSN NA forward primer: 5’-ACA ACG GCA TAA TAA CTG AAA CC-3’; WSN NA reverse primer: 5’-CAG GTA CAT TCA GAC TCT TGT GTT-3’. The relative viral copy number quantification was calculated from the slope of a standard curve that was obtained by using serial dilution of the corresponding plasmid as template.

### Fluorescence *In Situ* Hybridization

One day before the experiment, MDCK-SIAT1 cells were seeded on coverslips in 24-well plates. Cells were infected by viruses at an MOI of 5. Infection was synchronized and secondary infection was blocked using 20 mM NH_4_CI. Coverslips with cells were collected and washed twice with PBS, followed by fixation with 4% formaldehyde for overnight at 4 °C. To visualize viral RNA, RNA fluorescence *in situ* hybridization (RNA-FISH) through hybridization chain reaction (HCR)^63^ was then performed according to the manufacturer’s instructions with modifications. Cells were washed with PBS thrice, then permeabilized with 0.1% v/v Triton X-100 for 10 min at room temperature. After the permeabilization, cells were washed twice with 2× sodium chloride sodium citrate (SSC) buffer and incubated in HCR hybridization buffer at 37 °C for 30 min with gentle rocking. Subsequently, cells were incubated with hybridization buffer containing 1.2 pmol of the PB2 vRNA probe at 37 °C for overnight with gentle rocking.

The next day, excess probes were removed by washing cells 4 times with probe wash buffer at 37 °C with gentle rocking and were further washed twice with 5× SSC-T buffer (5× SSC with 0.1% Tween-20) for 5 min each at room temperature. Cells were then incubated with amplification buffer for 30 min at room temperature with gentle rocking. 18 pmol of B2-Alexa 647 amplifier hairpins per sample were snap-cooled by heating at 95 °C for 90 s and cooled to room temperature in the dark for 30 min. Hairpin solution, which was prepared by adding amplifier hairpins to 300 μL amplification buffer per sample, was added to samples and incubated for 1 h at room temperature in the dark with gentle rocking. Excess hairpins were removed by washing samples 5 times with 5× SSC-T buffer. For the first wash, samples were incubated in 5× SSC-T buffer supplemented with 1 μg/mL DAPI (Invitrogen) for 20 min at room temperature with gentle rocking. All subsequent washes were performed for 5 min at room temperature with gentle rocking. After the last wash, samples were mounted on glass slides with ProLong Diamond Antifade Mountant (Invitrogen). Cells were visualized using a LSM880 microscope system (Zeiss).

### Micrograph Analysis

Micrographs were analyzed using a custom pipeline on CellProfiler v4.2.1^64^ (Broad Institute). Using images from the DAPI channel, nuclei were identified using a minimum cross-entropy thresholding method. Then, nuclei were used to propagate and identify cells using the Otsu thresholding method. Nuclei and cells that touch the border of the image were discarded. Cytoplasm was identified by subtracting nuclei from cells. The ratio of the mean intensity of PB2 vRNA in the nucleus to that of PB2 vRNA in the cytoplasm was subsequently calculated. Only cells with PB2 vRNA median intensity of at least 0.05 were considered.

### Western blot analysis

Cells were lysed in RIPA lysis buffer and the cell pellets were removed by centrifugation at speed of 14,000 rpm for 30 min at 4 °C. Protein samples were prepared by mixing with 4× Laemmli Sample Buffer (Bio-Rad) supplemented with β-mercaptoethanol (Sigma-Aldrich) and boiled at 95 °C for 10 min. After SDS-PAGE, the proteins were transferred from the gel to polyvinylidene fluoride membranes (Bio-Rad). Membranes were blocked with 5% skim milk in PBST (PBS supplemented with 0.1% Tween-20) for 1 h at room temperature and incubated overnight with primary antibodies diluted in 5% skim milk in PBST at 4 °C. Membranes were washed three times, 10 min each with PBST, incubated with secondary antibodies for one hour at room temperature. Afterwards, membranes were washed three times, 10 min each with PBST. Positive immunostaining bands on the membranes were visualized using ECL Select Western Blotting Detection Reagent (Cytiva) and scanned by iBright 1500 imaging system (Invitrogen).

### Growth kinetic analysis of virus

Confluent MDCK-SIAT1 or A549-Dual cells seeded in 24-well plates were washed with PBS once and infected with virus at the indicated MOI. After 1 h of adsorption, the viral inoculum was removed, and infected cells were washed twice with PBS, and then cultured in DMEM medium supplemented with either 0.5 μg/ml TPCK-treated trypsin for A549 or 1 μg/ml TPCK-treated trypsin for MDCK-SIAT1 at 37 °C. Supernatants were collected at the indicated time points by centrifugation at 13000 × g for 1 min to remove dead cells and stored at −80 °C until being titrated. Virus titers were determined by TCID_50_ assay in MDCK-SIAT1 cells.

### Quanti-luc luciferase assay

QUANTI-Luc Gold is a two-component reporter kit which contains: QUANTI-Luc Plus and QLC Stabilizer (Invivogen). A standard protocol according to manufacturer’s instructions was followed. Quanti-Luc pouches were dissolved in sterile water together with QLC stabilizer. 20 μL of cell supernatant was added to a white opaque 96-well plate. 50 μL of QUANTI-Luc Gold assay solution was added to each well. The measurement was carried out immediately using BioTek synergy HTX multimode reader (Agilent).

### Quanti-blue SEAP phosphatase assay

QUANTI-blue Gold is a two-component kit which contains: QB reagent and QB buffer (Invivogen). A standard protocol according to manufacturer’s instructions was followed. The following protocol refers to the use of 96-well plates. 180 μL of QUANTI-Blue Solution was dispensed per well into a flat-bottom 96-well plate. 20 μL of sample (supernatant of SEAP-expressing cells) or negative control (cell culture medium) were added and incubated at 37 °C for 6 h. Optical density (OD) at 620 nm was measured using a BioTek synergy HTX multimode reader (Agilent).

### ISRE luciferase assay

For luciferase assays, HEK293T cells were seeded in a 24-well plate at a density of 100,000 cells per well. The next day, cells were transiently transfected with pRL-TK and ISRE-Luc reporter plasmids along with the indicated plasmids for 24 h. Cells were then treated overnight with 2000 U/mL of universal Type I IFN (PBL Bioscience) and lysed at 24 h post treatment using Passive Lysis Buffer (Promega). Samples were processed and luciferase activity was measured using the Dual-Luciferase Assay System (Promega) according to the manufacturer’s instructions. Measurements were acquired by BioTek synergy HTX multimode reader (Agilent). Firefly luciferase values were normalized to Renilla luciferase values.

### Caspase-3/7 activation assay

Cells were stained with CellEventTM caspase-3/7 green flow cytometry assay kit (Thermo, catalog #: C10427) according to the manufacturer’s protocol. Data were acquired by a BD Symphony A1 (BD Bioscience) flow cytometer, and the results were analyzed in FlowJo v10.8 software (BD Life Sciences).

### Influenza virus polymerase activity assay

Dual luciferase activity reporter assays were performed to compare the polymerase activity of RNP complexes in the presence of indicated NEP plasmid. RNP complex expression plasmids composed of PB2, PB1, PA (125 ng each) and NP (250 ng), together with pYH-Luci reporter plasmid (125 ng) and Renilla reporter plasmid (10 ng) were co-transfected into HEK293T cells. At 24 h after transfection, cells were lysed, and the luciferase activity was measured using the Dual-Luciferase reporter assay system (Promega). Measurements were acquired by a BioTek synergy HTX multimode reader (Agilent).

### Quantification and statistical analysis

All the statistical analyses have been performed using Prism 9 Graph Pad Software. Two-tailed Student’s unpaired t-test was performed to compare between two populations of data (e.g., control and sample) whereas two-way ANOVA was applied for multi sample comparisons. All data generated were from independent biological replicates where n ≥ 3, each measured in technical duplicates or triplicates. Results have been presented as means ± standard deviation (SD) or standard error of the mean (SEM).

### Data availability

Raw sequencing data have been submitted to the NIH Short Read Archive under accession numbers: BioProject PRJNA1083715 and BioProject PRJNA1110270.

### Code availability

Custom python scripts for all analyses have been deposited to: https://github.com/nicwulab/WSN_NEP_DMS.

